# The impact of *Piscirickettsia Salmonis* infection on genome-wide DNA methylation profile in Atlantic Salmon

**DOI:** 10.1101/2021.12.20.473279

**Authors:** Robert Mukiibi, Carolina Peñaloza, Alejandro Gutierrez, José M. Yáñez, Ross D. Houston, Diego Robledo

**Affiliations:** The Roslin Institute and Royal (Dick) School of Veterinary Sciences, The University of Edinburgh, Edinburgh, UK; Institute of Aquaculture, Faculty of Natural Sciences, University of Stirling, Stirling FK9 4LA, UK; Facultad de Ciencias Veterinarias y Pecuarias, Universidad de Chile, Santiago, Chile; Center for Research and Innovation in Aquaculture (CRIA), Universidad de Chile, Santiago, Chile

**Keywords:** *Salmo salar*, aquaculture, RRBS, disease resistance, bacteria

## Abstract

Salmon rickettsial septicaemia (SRS), caused by the intracellular bacteria *Piscirickettsia Salmonis*, generates significant mortalities to farmed Atlantic salmon, particularly in Chile. Due to its economic importance, a wealth of research has focussed on the biological mechanisms underlying pathogenicity of *P. salmonis*, the host response, and genetic variation in host resistance. DNA methylation is a fundamental epigenetic mechanism that influences almost every biological process via the regulation of gene expression and plays a key role in the response of an organism to stimuli. In the current study, the role of head kidney and liver DNA methylation in the response to *P. salmonis* infection was investigated in a commercial Atlantic salmon population. A total of 66 salmon were profiled using reduced representation bisulphite sequencing (RRBS), with head kidney and liver methylomes compared between infected animals (3 and 9 days post infection) and uninfected controls. These included groups of salmon with divergent (high or low) breeding values for resistance to *P. salmonis* infection, to examine the influence of genetic resistance. Head kidney and liver showed organ-specific global methylation patterns, but with similar distribution of methylation across gene features. Integration of methylation with RNA-Seq data revealed that methylation levels predominantly showed a negative correlation with gene expression, although positive correlations were also observed. Methylation within the first exon showed the strongest negative correlation with gene expression. A total of 911 and 813 differentially methylated CpG sites were identified between infected and control samples in the head kidney at 3 and 9 days respectively, whereas only 30 and 44 sites were differentially methylated in the liver. Differential methylation in the head kidney was associated with immunological processes such as actin cytoskeleton regulation, phagocytosis, endocytosis and pathogen associated pattern receptor signaling. We also identified 113 and 48 differentially methylated sites between resistant and susceptible fish in the head kidney and liver respectively. Our results contribute to the growing understanding of the role of methylation in regulation of gene expression and response to infectious diseases, and in particular reveal key immunological functions regulated by methylation in Atlantic salmon in response to *P. salmonis.*

## 1. Introduction

DNA methylation is a fundamental epigenetic mechanism that involves the conversion of cytosine (predominantly in the CG or CpG dinucleotides) to 5’-methylcytosine by DNA methyltransferases (DNMTs) such as *DNMT3A* and *DNMT3B* [1]. DNA methylation plays a significant role in maintaining the stability of the genome by silencing transcription of repetitive elements in the genome [1]. While DNMTs promote methylation, ten eleven translocation (TET) methylcytosine dioxygenases including *TET1, TET2* and *TET3* promote demethylation of the methylated cytosines [2, 3]. It is the interplay of DNMT and TETs (i.e. methylation and demethylation) that transcends into coordinated transcriptome changes to ensure appropriate cellular response to internal or external stimuli such as pathogen infection [4, 5]. Together with other epigenetic mechanisms, DNA methylation is involved in transcriptional gene expression regulation, and thus it is involved in modulating crucial biological processes including cell differentiation, proliferation, development and function of different tissues [6–8]. Increased methylation levels, especially in the promoter regions of genes, has been associated with repressed gene expression, however augmentation of gene expression by increased methylation has also been reported [5].

Pathogen infection can trigger epigenetic changes including DNA methylations that consequently lead to increased expression of immune genes in the host cells to combat and eradicate the intruding pathogen from the body [5, 6, 9]. Through host-pathogen co-evolution, pathogens have also developed mechanisms of manipulating their host’s methylomes through which they can regulate their host’s transcriptome (to dysregulate antipathogenic host genes or pathways) to enable their survival and multiplication within the cells [5, 10–12]. Elucidation of global or targeted DNA methylation changes upon *in vivo* and *in vitro* vertebrate infection by different pathogenic bacterial species such as *Mycobacteria tuberculosis* [12–14], *Escherichia coli* [15, 16] and *Salmonella* spp [17, 18] has been demonstrated in different species. Some of these methylation changes have been correlated with crucial genes in the innate and adaptive immune systems of the host. It has been suggested that identification of methylation signatures associated with pathogen infection could aid in the development of specific therapeutics [5, 6], diagnostic or biomarker tools [5] and vaccines [19]. Additionally, for livestock and aquaculture species, identification of DNA methylation signatures related with genetic variation in host resistance enables identification of functional and regulatory genomic features that could be integrated into genomic selection to sustainably improve resistance to the infection [20, 21].

*Piscirickettsia salmonis* (*P. salmonis*) is a major pathogenic threat to farmed salmonid fish industry by causing salmon rickettsial septicaemia (SRS) disease*. P. salmonis* is a gram negative facultative intracellular bacteria that mainly invades and multiply in the macrophage cells [22]. SRS has been reported in numerous farmed salmon producing countries including Chile, Norway, Scotland, Canada, and the United States [23]. In Chile, where this disease is endemic, it accounted for more than 50% of the total farmed Atlantic salmon infectious disease-related mortalities in the first semester of 2020 and 2021[24]. SRS causes significant mortalities and morbidity in the seawater stage of the production cycle, and results in annual loses of US$300M – US$700M, hence making it one of the biggest problems faced by the Chilean salmon aquaculture industry [25–27]. To date, several strategies have been applied to prevent or control the disease, including vaccination, reduced stocking densities, antimicrobial treatments and increased biosecurity at farms [27, 28]. However, SRS continues to be a pressing industry problem [26–28], and the excessive use of antimicrobials can promote the evolution of antimicrobial resistance in bacterial communities [23, 26, 27, 29]. Genetic improvement of host resistance is a promising and complementary prevention strategy, and additive genetic variation for resistance to SRS has been detected in various farmed populations, yielding heritability estimates ranging from 0.11 to 0.43 [30–32]. In fact, genetic improvement for SRS resistance is already being implemented in different commercial breeding programs for Atlantic salmon in Chile [33], accompanied with incorporation of genomic information to increase accuracy on the estimation of the genetic merit of breeders[34, 35].

Significant effort has also been placed on understanding the host gene expression response to *P. salmonis* infection in Atlantic salmon, with some of the results suggesting that the pathogen modulates several aspects of the host immune response to facilitate its survival and replication [31, 36–40]. Epigenetic changes, such as DNA methylation, may potentially underlie these transcriptional modifications, and could play a role in the mechanisms of host genetic resistance. In this line, a study of Pacific salmon (*Oncorhynchus kisutch*) response to SRS highlighted variation in spleen methylation profile following infection, and linked this variation to immune response pathways [41]. However, the role of DNA methylation in response to SRS in Atlantic salmon, and its relationship to genetic variation in host response has not been studied. Herein, the global patterns of methylation in two Atlantic salmon lymphoid tissues (head kidney and liver) were examined following experimental infection with *P. salmonis*, and differences in methylation profile were examined between infected fish and controls, and also between fish with differential genetic resistance to SRS.

## 2. Materials and Methods

### 2.1. Animals and SRS challenge trial

A total of 2,265 Atlantic salmon pre-smolts (average weight 135 ± 47 g) from 96 full sibling families from the breeding population of AquaInnovo (Salmones Chaicas, X^th^ Region, Chile) were experimentally challenged with *P. salmonis* (strain LF-89) with random distribution across 3 x 7 m^3^ tanks. All fish were vaccinated against *Flavobacterium*, infectious pancreatic necrosis virus (IPNV; Alpha Ject Flavo + IPN) and infectious salmon anaemia virus (ISAV; Alpha Ject Micro 1-ISA). Prior to the challenge, animals were tested negative for for ISAV, IPNV, *Renibacterium salmoninarum, Flavobacterium psycrophilum* and Mycoplasma by qPCR, and for bacterial contamination using culture in TSA, TSA+salt, and *Piscirickettsia salmonis* agar at 18°C and 35°C. The fish were intraperitoneally injected with 0.2 mL of a 1/2030 dilution of *P. salmonis.* This dose of the inoculum was previously determined using a group of 300 fish from the same families that were challenged with different doses of the bacteria. It was expected that the dose would cause mortality of approximately 50% of the challenged fish. The inoculated fish were retained in the 3 x 7 m^3^ tanks (~755 animals per tank) throughout the trial, and the experiment was terminated after 45 days after mortality had plateaued. The physic-chemical conditions during the trial included; temperature 14 ± 0.0 °C, salinity 30.4 ± 0.5 %, pH 7.4 ± 0.1, and oxygen saturation 102.2 ± 6.0 %. All animals were genotyped and their estimated breeding values (EBVs) for resistance to SRS (measured as mortality / survival) were obtained for every animal (details are given in Moraleda et al. 2021). The EBVs were used to classify the animals into resistant and susceptible groups based on their estimated breeding values to enable differential methylation comparisons related to genetic resistance (see below).

The challenge experiments were performed under local and national regulatory systems and were approved by the Animal Bioethics Committee (ABC) of the Faculty of Veterinary and Animal Sciences of the University of Chile (Santiago, Chile), Certificate N° 01-2016, which based its decision on the Council for International Organizations of Medical Sciences (CIOMS) standards, in accordance with the Chilean standard NCh-324-2011.

### 2.2. Tissue collection

During the trial a subset of animals were sampled for transcriptomic (RNA-Seq) and methylation (Reduced Representation Bisulphite Sequencing, RRBS) analyses. All sampled animals were ethically anesthetized and euthanized before collecting the samples. In total, liver and head kidney tissue samples were collected from 144 animals at three time points: 48 unchallenged (control) fish, 48 pathogen-challenged fish at 3 days post infection (dpi), and 48 pathogen-challenged fish at 9 dpi. Tissues dissected from each animal were individually stored in RNAlater at 4°C for 24 hours, and thereafter kept at −20°C until DNA extraction, and RNA extraction as previously described by Moraleda *et al.* 2021.

### 2.3. RRBS library preparation and sequencing

A total of 66 samples (i.e. 33 samples of each tissue) including 7 x control, 13 x 3dpi, and 13 x 9dpi samples were selected for the RRBS analyses based on the availability of RNA sequencing data (from Moraleda *et al.* 2021), their EBVs for resistance to SRS, and the availability of high quality DNA for bisulfite conversion and sequencing. Bisulfite converted reduced representation genomic DNA libraries were prepared using the Diagenode Premium RRBS kit [42] following the manufacturer’s instructions. Briefly, 100 ng of genomic DNA from each of the 66 samples was digested with the restriction enzyme MspI for 12 hours, followed by fragment end-repair, A-tailing, and adapter ligation. Methylated and unmethylated spike-in controls were added to monitor bisulfite conversion efficiency. Individual libraries were quantified in duplicate by qPCR. Samples with similar qPCR threshold cycle (Ct) values were multiplexed in equimolar amounts in pools of six. Pools were then subjected to bisulfite conversion. Thereafter, RRBS libraries were enriched by PCR and purified with AMPure® XP beads (#A63881, Beckman Coulter). Quality assessment of the RRBS libraries was performed by verifying the fragment size distribution on an Agilent 2200 Bioanalyzer (Agilent Technologies). Libraries were quantified using a high sensitivity assay on a Qubit 3.0 Fluorometer (Life Technologies, ThermoFisher Scientific), and then pooled at equimolar concentrations for sequencing on three flow cell lanes of an Illumina NovaSeq S1 platform (50 bp paired-end sequencing) at Edinburgh Genomics (University of Edinburgh).

### 2.4. RRBS data processing and methylation profiling

Raw sequence read data quality was initially evaluated using FastQC software Version 0.11 (https://www.bioinformatics.babraham.ac.uk/projects/fastqc/). The raw sequence data were then cleaned by removing low quality base calls (Phred score < 20) from the ends of the reads, short reads (< 20bp) and Illumina sequencing adapters using Trim Galore Version 0.5.0 software [43] with default RRBS paired-end parameters. Methylation profiling from the cleaned sequence data was performed using Bismark pipeline tools Version 0.20.0 [44]. To facilitate bisulphite alignment, the Atlantic salmon reference genome (GCF_000233375.1_ICSASG_v2) was bisulphite converted *in silico* to C-> T (forward) and G-> A (reverse) versions using the bismark_genome_preparation script. The bisulphite-converted clean RRBS sequence reads were then aligned to the *in-silico* bisulphite-converted reference genome versions using the Bismark script that utilizes bowtie2 [45] as the underlying short read aligner. Subsequently, the methylation state of each cytosine in the genome was called from the alignments using the same Bismark script. The methylation call for all profiled cytosine (C) nucleotides were extracted from the alignment bam files using the Bismark methylation extractor script into CpG methylation coverage (count) files that were used for downstream analyses. Due to the inability of RRBS to distinguish between C/T single nucleotide polymorphisms (SNPs) and true C-T bisulphite conversion [46], all the C/T SNPs were filtered out from profiled CpG sites based on previous whole genome sequencing data from the same population [47].

### 2.5. Genomic annotation of CpG sites and functional enrichment analyses

The identified CpG sites were functionally annotated according to the genomic features in the salmon genome annotation file (https://ftp.ncbi.nlm.nih.gov/genomes/all/GCF/000/233/375/GCF_000233375.1_ICSASG_v2/GCF_000233375.1_ICSASG_v2_genomic.gtf.gz) using the annotatePeaks.pl tool from the HOMER software (http://homer.ucsd.edu/homer/ngs/annotation.html). CpG sites were annotated as located either in the putative promoters / transcription start site (TSS, defined as −1kbp to +100bp around the TSS), exons, introns, the transcription termination site (TTS, defined as −100bp to +1kbp around the TTS) or in intergenic regions.

### 2.6. Differential methylation analyses

Differential methylation analysis was performed using a Bioconductor R package edgeR Version 3.28.1 [48]. Firstly, CpG sites which had low coverage (< 10 reads per sample), those that were either always methylated or unmethylated across all samples, and those located on unplaced-scaffolds without annotated genes were removed from the analysis. Principal component analysis was performed on the methylation proportion with the *prcomp* function implemented in R to visualise the distribution of the samples according to their overall methylation patterns, and outlier samples notably isolated from the major clusters were discarded. 6 control, 12 x 3 dpi and 12 x 9 dpi samples were retained for differential methylation analysis in the liver, and 7 control, 11 x 3 dpi and 9 x 9 dpi samples were retained for analysis in the head kidney. The read counts of each CpG site within a sample library were then normalized by scaling both methylated and unmethylated counts to the average library size of the sample (i.e., average of methylated and unmethylated libraries). A negative binomial generalized linear model implemented in edgeR [48] was used to model the normalized counts for each infection stage (control, 3 dpi and 9 dpi) of the animals. Differential methylation between sample groups was tested via likelihood ratio tests (LRT), and a CpG site was considered significantly differentially methylated with a Benjamini - Hochberg corrected P-value < 0.1. Additionally, the samples were classified into resistant or susceptible groups based on genomic breeding values [31], with 11 resistant and 9 susceptible samples for liver, and 7 resistant and 12 susceptible samples for head kidney; resistant and susceptible samples were analysed for differential methylation within each group (control, 3dpi, 9dpi). The R package Circulize Version 0.4.10 [49] was used to visualize genome-wide methylation patterns and differences in methylation between groups. Those CpG sites located in or in close proximity to genes as described above were used for functional pathway enrichment analyses. Pathway enrichment analysis was performed using the KEGG Orthology-Based Annotation System (KOBAS) Version 3.0.3 [50], pathways showing a p-value < 0.05 were considered significantly enriched.

### 2.7. Integrating DNA methylation and transcriptome expression

Whole transcriptome gene expression data (counts per million or cpm) for the samples used in the current study were obtained from Moraleda *et al* [31]. For each tissue, Pearson’s correlations between DNA methylation proportion of the CpG sites and cpm of their respective annotation gene transcripts were computed for each methylation site – gene expression pair using the Hmisc [51] package in R. Pearson’s correlations between average (across samples) transcript expression and average DNA methylation within the different annotation genomic features (i.e., promoter-TSS, introns, exons, TTS, and intergenic regions as defined above) were also computed.

## 3. RESULTS

### 3.1. Sequencing and DNA methylation summary statistics

Sequencing of the liver and head kidney RRBS libraries yielded on average 40.1 ± 2.2 and 41.2 ± 1.87 million (mean ± SD) raw paired-end reads after quality control, respectively. There were no significant differences between head kidney and liver samples (Figure S1A in Supplementary file 1). An average of 42.5% of the reads aligned to a unique position in the Atlantic salmon genome assembly (Figure S1B in Supplementary file 1). Most of the methylated cytosines identified in the sequenced libraries were within CpG sites (>84%; Figure S1B and Figure S1C in Supplementary file 1), and on average 3.6 ± 0.1 million and 3.8 ± 0.05 million CpG sites were profiled per sample in the liver and head kidney, respectively (Figure S1D in Supplementary file 1). The methylation levels of 693,215 and 961,595 CpG sites were profiled in all liver and head kidney samples respectively.

### 3.2. Atlantic salmon head kidney and liver methylation patterns in unchallenged fish

A total of 308,198 and 247,252 CpG sites with sufficient sequencing coverage (≥10 read counts) were identified as showing inter-individual methylation variation across head kidney and liver control samples respectively. Of these variably methylated sites, 10,474 sites across the genome were significantly differentially methylated between the two tissues, 6,456 hyper-methylated and 4,018 hypo-methylated in the head kidney samples relative to the liver samples (Figure 1A). Additionally, 8719 sites were identified as fully unmethylated across all head kidney samples, while 10,743 sites were identified as fully unmethylated across all liver samples. These sites generally had differing density distribution patterns across the genome in the two tissues (Figure 1A). 1,446 and 19,741 CpG loci were identified as fully methylated across all head kidney and liver samples respectively. In both liver and head kidney, both fully methylated and variably methylated CpGs were predominantly located in intergenic and intronic regions (>85% of all methylated sites), while fully unmethylated sites were predominantly located in promoter-TSS and exonic regions (>60% of all unmethylated sites) (Figure 1B). For the CpG sites showing variable levels of methylation within each tissue, they were mainly located in the intergenic (49.9 ± 0.94%) and intronic (39.0 ± 0.49%) regions (Figure 1B). Additionally, variable methylation levels showed a bimodal distribution (Figure 1C). While intragenic regions (exons and introns), TTS, and intergenic regions were heavily skewed towards high (>75%) methylation (84.6 ± 2.45% of the CpGs), in promoter-TSS regions there was a higher proportion (39.3 ± 1.28% of the CpGs) of low (<25%) methylation levels (Figure 1C). Infection with *P. salmonis* did not significantly modify the distribution of methylation patterns described above (Figure S2 in Supplementary file 1).

**Figure 1:**
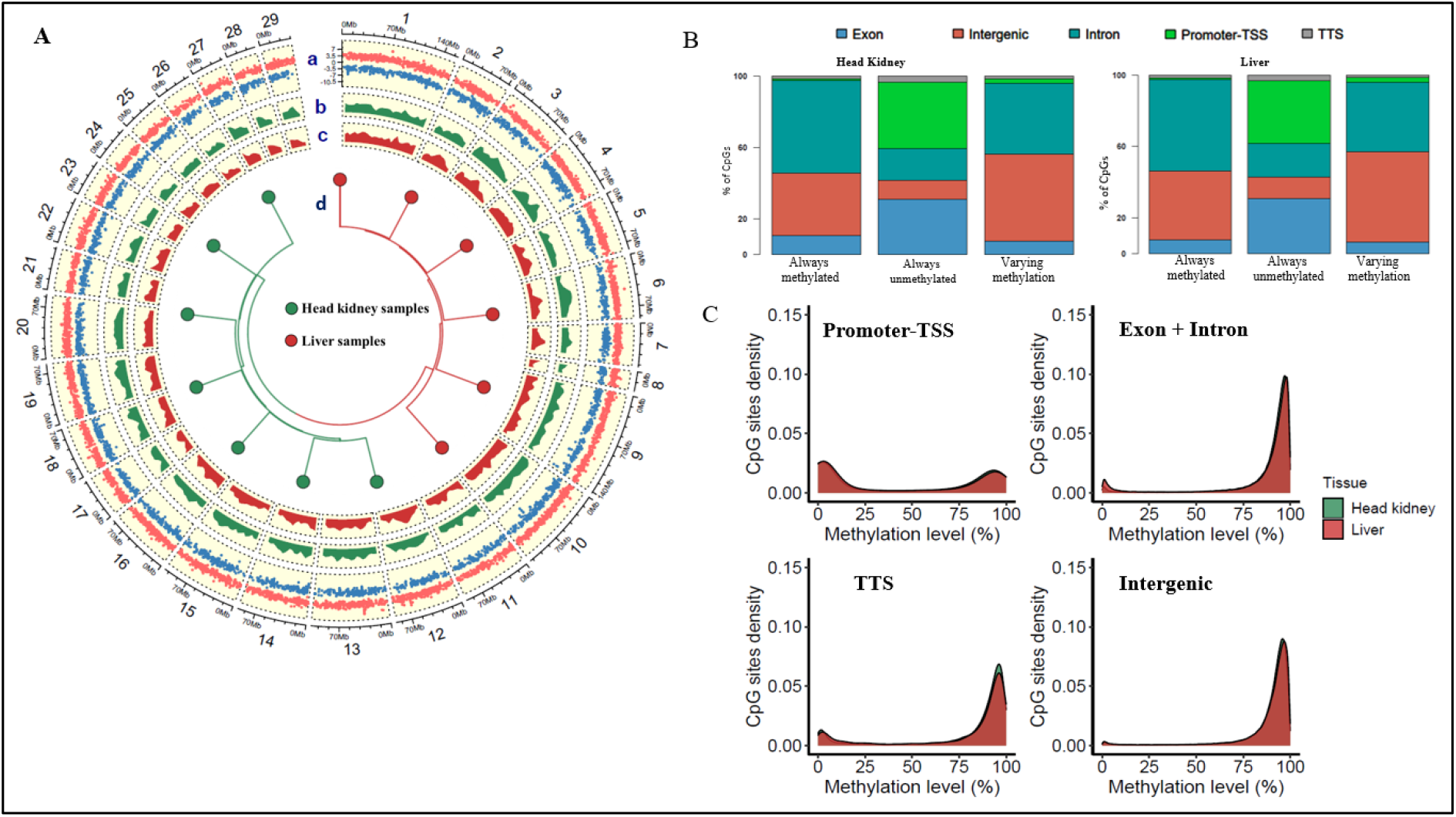
General methylation patterns in head kidney and liver tissue of unchallenged animals. A) Circos plot showing differential methylation between head kidney and liver samples (Track: a), Genomic distribution of always fully unmethylated CpG sites in the head kidney tissue (Track: b), Genomic distribution of always fully unmethylated CpG sites in the head kidney tissue (Track: c), Dendrogram plot showing unchallenged fish head kidney and liver samples clustering based on their methylation profiles(Track: d). B) Distribution of the three classes of methylation (i.e., fully methylated in all samples, fully unmethylated in all samples and varying methylation between samples.) in the different genomic features. C) Methylation variability within the different genomic features for the head kidney and liver tissues.

### 3.3. Integrating gene expression and DNA methylation

To understand the potential impact of methylation on transcriptional regulation, the correlation between the level of methylation and gene expression was assessed. 2,310 and 1,306 methylation sites showed significant correlation with the expression of the genes they are located in or neighboring in head kidney and liver, respectively (Supplementary file 2). The highest correlations were >|0.7|, but most of the values ranged between |0.4| and |0.5| (Figure 2A and B). Approximately 60% of the methylated sites showed negative correlations with gene expression, while 40% showed positive correlation, and the sign of the correlation was independent of the genomic feature location of the CpG site (Supplementary file 2). Overall, average methylation level within the promoter-TSS regions, the 1^st^ exon, and the TTS region showed the strongest negative correlations with gene expression, and methylation in the first intron also showed a weaker but significant negative correlation (Figure 2C and D). Methylation of CpGs within the other exons, other introns and intergenic regions showed weaker effect on gene expression in the two tissues (Figure 2C and D).

**Figure 2:**
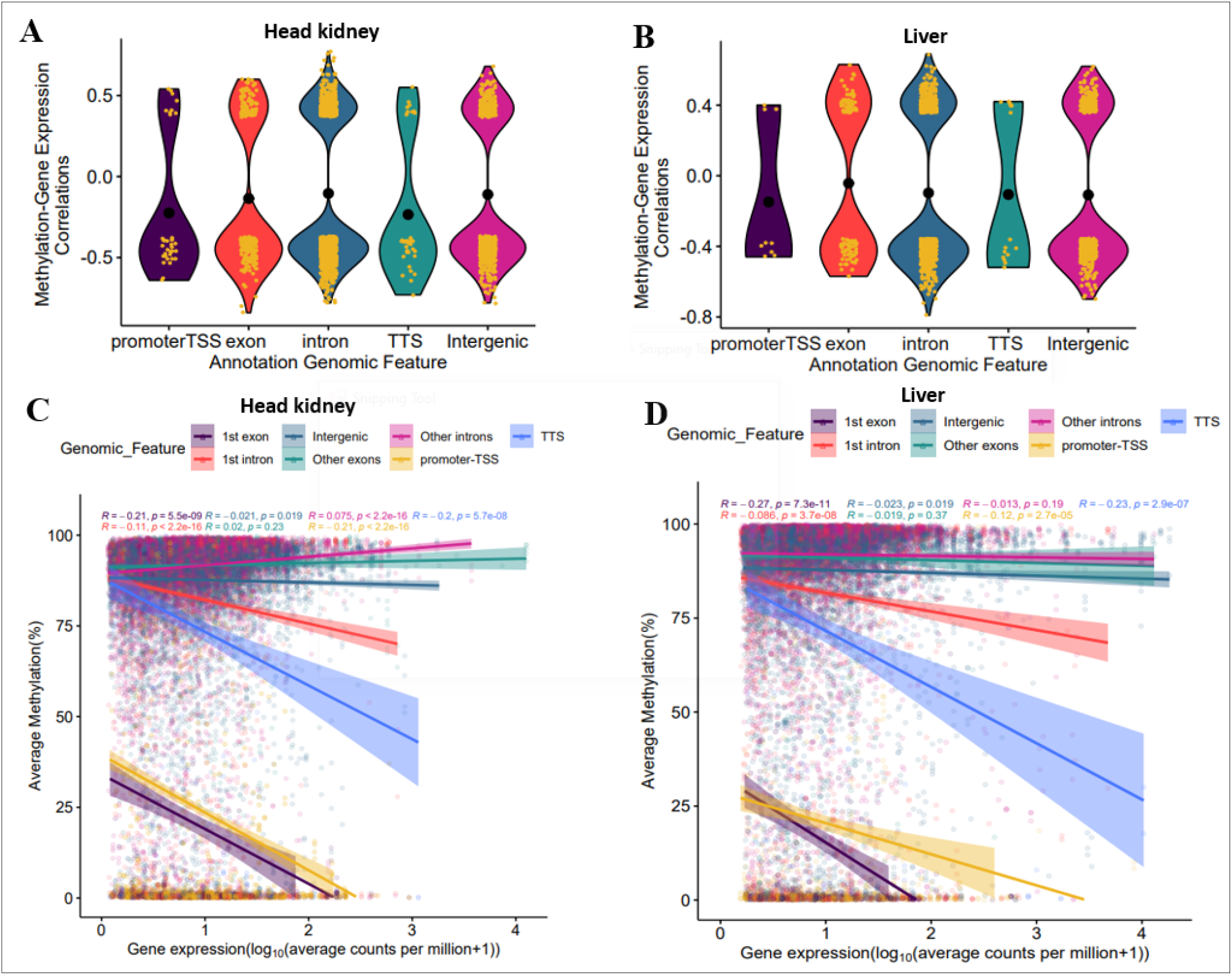
Global correlation between methylation and gene expression; A) Violin plot showing the distribution of magnitude of the correlations between DNA methylation and gene expression in head kidney, B) Violin plot showing the distribution of the magnitude distribution of the correlations between DNA methylation and gene expression in liver tissue, C) correlation plot showing general correlation between average gene expression and average methylation and in the different genomic features in the head kidney tissue, D) correlation plot showing general correlation between average gene expression and average methylation and in the different genomic features in the liver tissue.

### 3.4. SRS-induced changes in head kidney and liver methylation patterns

Comparison of the methylomes of *P. salmonis*-infected and healthy animals revealed a greater number of sites with significant genome-wide differences in methylation levels in head kidney than in liver samples (Figure 3A, Supplementary file 3). In head kidney, 911 CpG sites (DMCpGs) showed differential methylation between control and challenged animals at 3 day post infection (3 dpi), with 606 sites showing increased methylation levels (hypermethylation) upon infection and 306 showing decreased levels (hypomethylation) (Figure 3A, Supplementary file 3). At 9 days post infection (9 dpi) fewer significant differences were identified (813 DMCpGs), of which 405 were hyper-methylated and 408 hypo-methylated (Figure 3A) in the infected animals. In contrast, the liver just showed 30 DMCpGs between controls and 3 dpi samples (28 hyper-methylated and 2 hypo-methylated in infected samples), and 44 DMCpGs between controls and 9 dpi samples (36 hyper-methylated and 8 hypo-methylated in infected samples) (Figure 3A). The differences in methylation between 3 dpi and 9 dpi were negligible in both tissues, and the fold changes of the DMCpGs at 3 dpi and 9 dpi relative to control samples showed a high positive correlation (Figure 3B), suggesting that methylation patterns change following infection, but less so during the course of the infection between 3 and 9 dpi. This is also observed in the heatmaps showing the methylation patterns of the DMCpGs in both tissues (Figure 3C), where control and infected samples clustered separately with contrasting methylation patterns consistent across each group. This suggests that the changes in methylation are biologically relevant and play a role in the response to *P. salmonis* infection.

**Figure 3:**
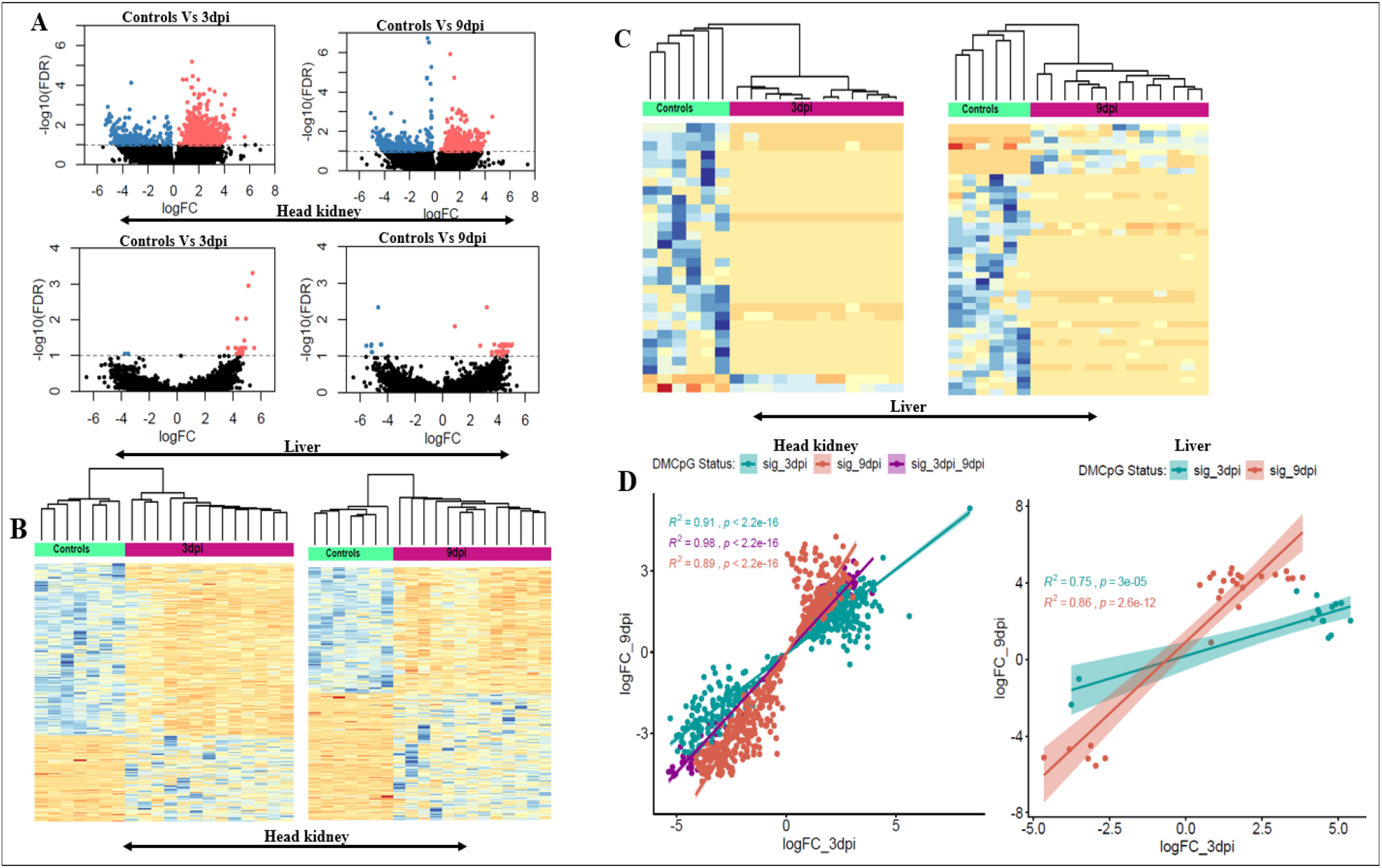
Differential methylation induced by P. salmonis infection; A) Volcano plots showing differential methylation between infected and unchallenged control animals at 3 and 9 days post infection. Significance at False discovery rate < 0.1. B) Heatmaps showing hierarchical clustering samples based on the differentially methylated sites. D) Correlation plots showing correlation between the difference in methylation at 3 days and 9days post infection in the two tissues.

### 3.5. Functional enrichment of DMCpGs induced by P. salmonis infection

The DMCpGs were assigned to genes according to their position in the annotated genome, and a functional enrichment analysis was performed for each tissue separately (Supplementary file 4). In head kidney, key pathways related to innate and adaptive immunity showed significant enrichment, including regulation of actin cytoskeleton (Figure 4), T cell receptor signaling pathway, bacterial invasion of epithelial cells, inflammatory mediator regulation of TRP channels, Fc gamma R-mediated phagocytosis and TNF signaling pathway. Several of these pathways were also identified in the gene expression analyses from the same dataset described by Moraleda et al. [31] and in previous genome-wide association studies (GWAS) for *P. salmonis* resistance in different salmonid species [52, 53], including the regulation of the actin cytoskeleton (Figure 4), which relates to the known role of *P. salmonis* to induce cytoskeletal reorganisation via actin depolymerisation [40]. While not significantly enriched, there were other pathways of interest with substantial numbers of differentially methylated genes, such as B cell receptor signaling pathway, phagosome, apoptosis, PI3K-Akt signaling pathway, Toll-like receptors signaling pathway, NOD-like receptor signalling pathway, cytokine-cytokine receptor interaction, lysosome, endocytosis, leucocyte transendothelial migration, tuberculosis and ubiquitin mediated proteolysis pathway (Supplementary file 3). Some of the key genes involved in multiple of these pathways included *PIK3R3, PIK3R1, ITGA5, VAV3,VAV2, AMPH, CLA, RAB5C, CBLB, NFATC1, NCF1, FADD, ROCK2* and RhoA, of which transcriptome expression of *PIK3R3* and *ITGA5* had significant negative correlation (−0.4 and −0.64 respectively) with DNA methylation.

**Figure 4:**
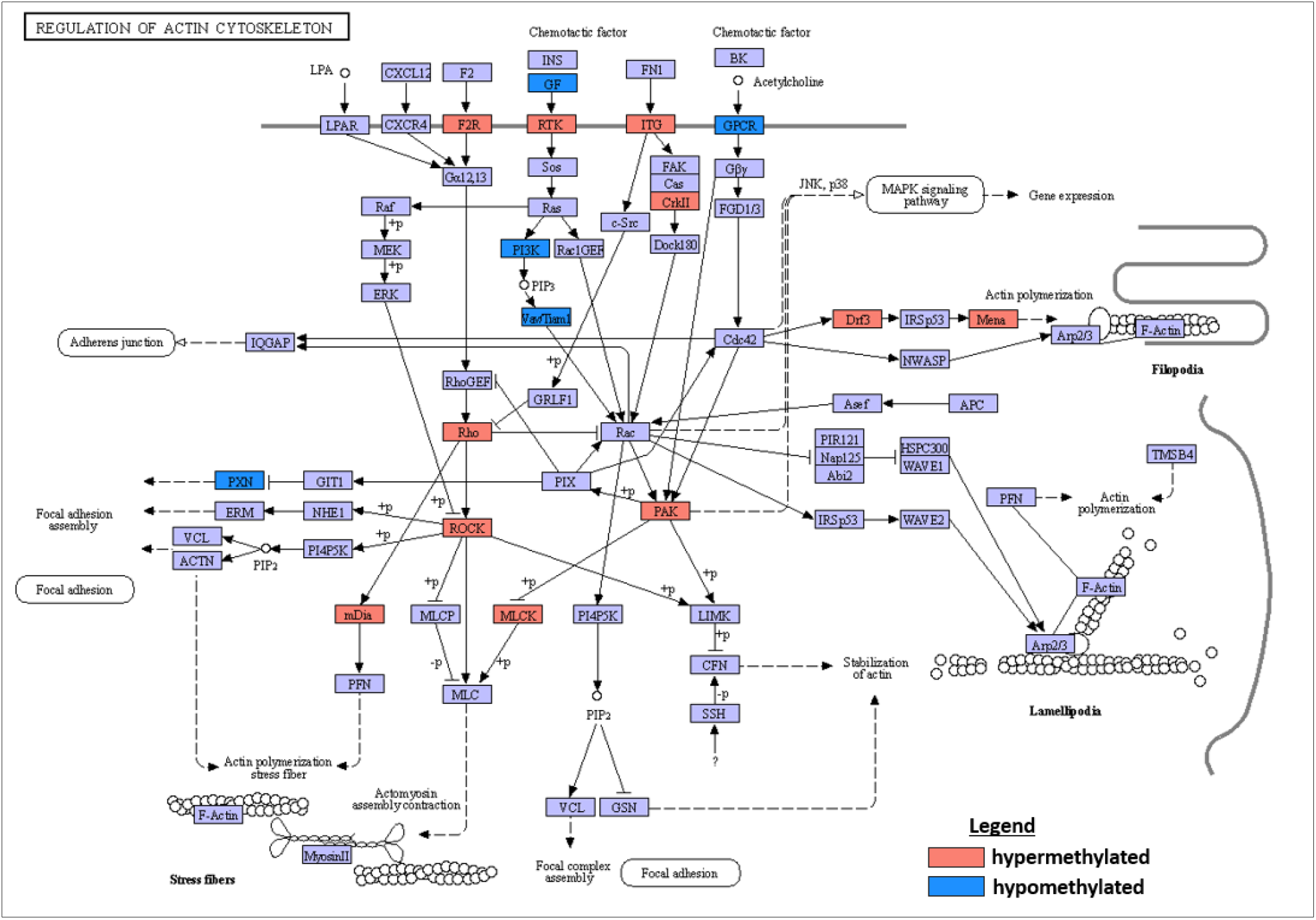
Regulation of actin cytoskeleton pathway. One of pathways enriched by genes showing significant altered methylation in response to P. salmonis infection in the head kidney.

### 3.6. Differential methylation between resistant and susceptible fish

Comparison of the methylation levels of *P. salmonis* resistant and susceptible fish (based on divergent EBVs) revealed 113 DMCpGs in the head kidney (63 hypomethylated and 50 hyper-methylated in resistant fish) and 48 DMCpG sites in the liver (23 hypo-methylated and 25 hyper-methylated in resistant fish) (Figure 5A, Supplementary file 5). The trends of methylation for the DMCpGs in each tissue were relatively consistent among susceptible animals or resistant groups (Figure 5B). Some of the DMCpGs identified in the head kidney tissue are located within or neighboring well-known immune related gene. These included *TUBA1A, CFL2* and *MTSS1* which are involved in cytoskeleton structure, and rearrangement regulation [54–56], and may relate to the previously identified enrichment of the actin cytoskeleton pathway. In addition, *CXCL12, CCR6, TNFRSF4* and *JAK2* were identified as differentially methylated between resistant and susceptible animals, and these are all related to cytokine signaling [57]. In the liver tissue, genes of interest that contained or were neighbouring DMCpGs included *DOCK1* and septin-2 both involved in cytoskeleton pathways [58, 59], and *BPI* which is involved in recognition and neutralization gram-negative bacteria lipopolysaccharide antigens [60].

**Figure 5:**
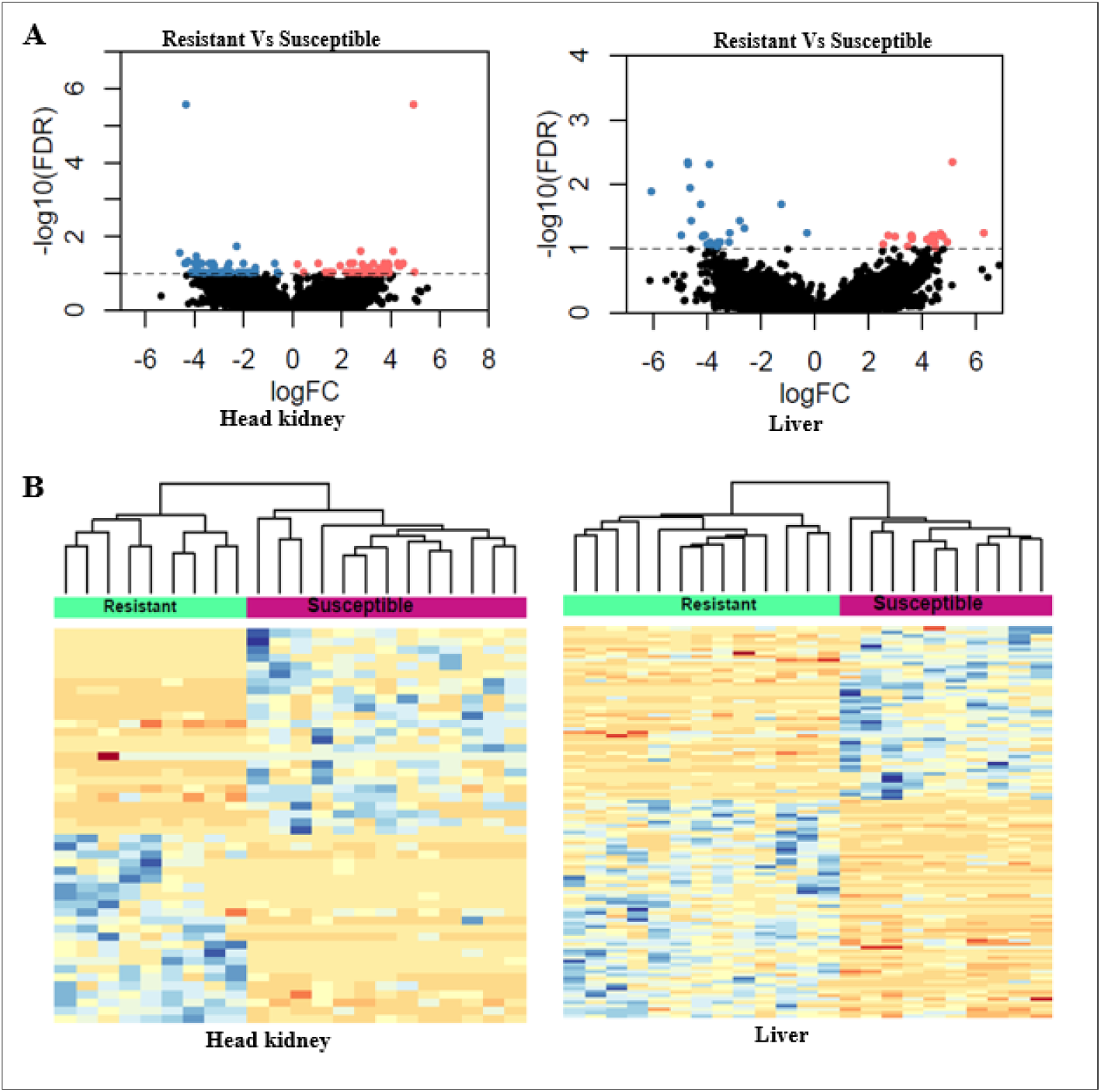
Differential methylation between resistant and susceptible animals based on EBVs; A) Volcano plots showing differential methylation the head kidney and liver tissue between resistant and susceptible animals, B) Heatmaps showing hierarchical clustering samples based on the differentially methylated sites.

## 4. Discussion

In the current study, the methylation patterns of head kidney and liver in Atlantic salmon and their remodelling in response to *P. salmonis* infection were investigated. Our study demonstrated significant genome wide methylation differences between the two tissues, although the methylation patterns within the different genomic features were conserved across tissues and conditions. Generally, methylation, especially in the promoter and at the start and the end of transcription regions, is negatively correlated with gene expression, although a significant number of sites showed positive correlation trends with gene expression. Infection by *P. salmonis* induces methylation changes in both tissues, with a larger impact on the head kidney methylome. The genes associated with these changes in methylation are involved in several key innate and acquired immune pathways, suggesting that DNA methylation plays a role in coordinating the Atlantic salmon immune response to *P. salmonis* infection.

The noticeable differences between the methylomes of head kidney and liver are a reflection of the methylation signatures that shape organ and cell differentiation during development [61], which allow for function specialization of the different tissues and organs. Differing methylation signatures between different tissues or cell types have previously been reported in mammalian [62, 63] and avian species [64]. In contrast, the distribution of methylation across genomic features is consistent between the two tissues. CpG sites in intragenic and intergenic regions showed high levels of methylation, whereas in promoter regions methylation showed more variation, with comparable numbers of highly and lowly methylated CpG sites. Similar methylation pattern have been observed in the gill tissue of Atlantic salmon [65], as well as in the testis, muscle, liver and spleen tissue of European sea bass [66]. The high methylation levels in both intragenic and intergenic regions is thought to contribute transcription repression of unwanted noncoding RNAs and repetitive genomic elements that are usually located within these regions, hence maintaining genome stability [67]. Hypermethylation within intragenic regions has also been demonstrated to repress the establishment of alternative transcription start sites that would lead to transcription of alternative transcripts [68], which could be deleterious or of no particular function in the cell [69]. The bimodal distribution observed in the promoter-TSS regions is consistent with the highly conserved architecture of promoters in vertebrate genomes, which contain two classes of promoters, one with high number of CpGs that are hypomethylated and another one with few CpGs that are hypermethylated [70]. The hypermethylated and hypomethylated promoters are associated with tissue-specific and broadly expressed genes respectively [70].

The strong negative correlation between gene expression and the methylation level of the 1^st^ exons observed in this study is consistent with previous observations in human cells [71, 72] and sea bass testis [66]. These observations demonstrate deviation from the canonically accepted mechanism which suggests that gene expression is regulated through modulation of DNA methylation within the promoter regions of genes [71]. The exact biological mechanisms responsible for the strong negative influence of 1^st^ exon methylation levels remain largely unknown. However, in vertebrates, CG content peaks at the 1^st^ exon /1^st^ intron junctions [73], hence making the 1^st^ exon a hotspot region for the recruitment of transcriptional factors to initiate transcription. Methylation at the transcription termination sites also showed a noticeably strong significant negative correlation with gene expression. We speculate this correlation could be due to potential overlaps between transcription sites and downstream enhancers [74, 75].

In any case, significant correlations between gene expression and methylation levels were observed for sites in all genomic features, indicating that methylation modulation of gene expression is not restricted to the promoter regions in the genome as classically regarded [71]. The correlations were predominantly (60%) negative, meaning that increasing methylation will generally results in a reduction of gene expression, in agreement with previous reports in fish and humans [66, 76]. However, many (40%) positive correlations between gene expression and methylation levels were also observed, implying that for a considerable number of genes increasing methylation would augment expression of these genes. These positive correlations between methylation and gene expression have been reported in multiple vertebrate studies [66, 76, 77], but the mechanism underlying these positive correlations remains poorly understood. It has been suggested that in some cases transcription can lead to DNA methylation, thus resulting into the positive correlation between methylation and gene expression [77].

Pathogen infections usually trigger host immune responses, which can lead to the activation of different immune cell types through changes in their gene expression [5, 6, 78]. DNA methylation is one of the well coordinated epigenetic mechanisms that contribute to the transcriptional reprogramming of host immune cells [5, 6, 78]. Interestingly, through millions of years of coevolution with their hosts, some pathogens have evolved the ability to modulate the expression of host immune genes via DNA methylation to promote their survival and multiplication in the host cells [78, 79]. In the current study, our results demonstrated that *P*. *salmonis* infection reshapes the head kidney methylome in Atlantic salmon, while inducing limited changes in the liver methylation profile. These results are consistent with the previous transcriptomic study on the same population used in this work, where a higher number of differentially expressed (DE) genes was observed in head kidney compared to the liver [31]. Nonetheless, Moraleda *et al* still reported substantial changes in the liver transcriptome in response to *P. salmonis* (>2000 DE genes) [31]. Therefore, while methylation potentially mediates an important part of the transcriptional changes, other mechanisms must be modulating the liver transcriptional response to the bacteria. This difference could be connected to the role of the head kidney, a primary lymphoid organ where immune cells go through differentiation and maturation [80, 81]. The head kidney is the main fish hematopoietic organ generating lymphoid and myelopoietic immune cells, key in antigen processing and antibody production, and also the main center for phagocytosis in the body, where destruction of pathogens including bacteria occurs [82, 83]. The differentially methylated sites found in the head kidney in response to *P. salmonis* in our study are located within or in close proximity to genes involved in immune pathways connected to the immune function of this organ.

The actin cytoskeleton modulates a wide range of immunological cellular processes or functions including phagocytosis, endocytosis, intercellular interaction, cell division, intercellular signal transduction, cell movement, activation, morphology and movement [58]. Therefore, alterations of the genes involved in the actin cytoskeleton can have significant impact on the immune system of the animal. Previously, Ramirez *et al* demonstrated that *P. salmonis* infection of Atlantic salmon macrophages disrupts cytoskeleton disorganization and increases the cells’ actin synthesis, which the bacteria utilize to generate vacuoles where they survive and multiply while shielding from cytosolic detection and destruction [40]. In agreement with these results, upon *P. salmonis* infection, we detected differential methylation of numerous (n = 40 gene transcripts) genes involved in the actin cytoskeleton regulation in the current study. These genes included four transmembrane receptors (*ITGA5, PAR1, CHRM3* and *PDGFRA)*, which are key in actin cytoskeleton signalling. Of these receptors, *ITGA5 (*integrin alpha-5) mRNA expression also had a strong negative correlation (−0.62) with methylation, and was previously identified as differentially expressed between *P. salmonis* infected and non-infected Atlantic salmon fish [31]. *PAR1, which* encodes proteinase-activated receptor 1, also had significant negative correlation (−0.59) between gene expression and methylation, and was also identified as differential expression between *P. salmonis* infected fish and healthy fish [31]. CrkII was another gene of interest involved in actin cytoskeleton regulation that showed differential methylation, significant negative correlation (−0.64) between gene expression and methylation, and was found to be differentially expressed between infected and unchallenged animals [31]. This gene encodes for the Crk adapter protein, which plays crucial roles in actin reorganization, phagocytosis, lymphocyte adhesion, activation and migration [84–86]. Previous GWAS for *P. salmonis* resistance in different salmonid species, including Atlantic salmon, coho salmon and rainbow trout, have also pinpointed actin cytoskeleton as an important mechanism of the host response to infection [52, 53]. Altogether, these results demonstrate the importance of both genetic variants and DNA methylation on the modulation of actin cytoskeleton in response to *P. salmonis* infection.

Precise pathogen recognition through pattern recognition receptors (PRRs) and associated pathways is important if the host organism is to establish a well coordinated, effective and rapid response against the pathogens [87]. Pathogen recognition triggers the initiation of the different immune response processes like phagocytosis, necrosis, production of type I interferons and inflammatory cytokines, and activation of the inflammasome in the phagocytic host cells [87, 88]. These processes directly serve in pathogen elimination, as well as recruitment and activation of more innate and adaptive immune cells to fight the infection [87]. In the current study, we identified some key genes with differential DNA methylation that are involved in cell transmembrane detection pathogenic bacteria via the Toll-like receptor (TLR) signaling pathway (*FADD, PIK3R3, PIK3R1* and *IL-12*) and intracellular bacteria recognition via NOD-like receptor signalling pathway (*Erbin* and *BIRC2). FADD* encodes for Fas-associated protein with death domain, a critical adaptor in the TLR activated apoptosis that oligomerizes caspase 8, which in turn activates other effector caspases to initiate apoptosis [89, 90]. *PIK3R3* and *PIK3R1* encode for phosphatidylinositol 3-kinases’ (PI3Ks) gamma and alpha regulatory subunits respectively [91]. PI3Ks modulate numerous immune response processes/pathways via the activation of serine/threonine-protein kinase (*AKT).* For instance, a previous GWAS study suggested PIK3AP1, a gene involved in B-cell development [92], as a strong candidate gene associated to *P. salmonis* resistance in coho salmon [93]. Within the TLR signaling pathway, PI3Ks are involved in modulating the synthesis of proinflammatory cytokines such as interleukin 12 (*IL-12*) by immune cells [94]. IL-12 is a key modulator of differentiation and maturation of Th1 cells from naive CD4+ T cells, proliferation of CD8+ T cells and Natural killer T (NKT) cells all of which play critical roles in the elimination of intracellular pathogens [94–96] such as *P. salmonis.* Previously Moraleda *et al.* reported differential expression of *IL-12* and *FADD* between *P. salmonis* infected and control fish from the same population used in the current study [31]. However, we did not observe significant correlation between methylation of these genes and mRNA expression in the present study.

Phagocytosis is major innate immune strategy through which pathogenic organisms (including bacteria) are destroyed or eliminated from the body by phagocytic cells [88]. Phagocytosis of bacteria is initiated upon detection of bacteria associated molecular patterns by pattern recognition receptors (PPRs) at the phagocyte cell membrane, or after detection of Ig-opsonized bacteria by the Fc receptors (FcR) on the cell surface [88, 97, 98]. Interestingly, in the current study, 18 genes with differential methylation are involved in the Fc gamma R-mediated phagocytosis pathway, including key genes such as *HCK, PIK3R3, PIK3R1, CrkII* and *NCF1. HCK* encodes for the hematopoietic cell kinase, an Src kinase that phosphorylates Fc receptors engaged with Ig-opsonised pathogens triggering actin cytoskeleton rearrangement, a key process in phagocytic pathogen engulfment [85]. As aforementioned *PIK3R3 and PIK3R1* encode for regulatory subunits of *PI3Ks. PI3Ks* are key enzymes that modulate cytoskeleton reorganization and formation of the phagosome via the PI3K-Akt signaling pathway [99]. Although the PI3K-Akt signaling pathway was not statistically significantly enriched in our study, 42 candidate genes involved in this pathway showed differential methylation, indicating its importance in the response to *P. salmonis* infection. Leiva *et al* reported significant differential methylation of genes within PI3K-Akt signaling pathway in the spleen tissue of Pacific salmon (*Oncorhynchus kisutch*) in response to *P. salmonis* infection [41]. Further, Moraleda *et al* identified multiple phosphatidylinositol 3-kinases and phosphatidylinositol 3-kinase regulatory subunit proteins as differentially expressed between infected and healthy animals [31]. *CrkII*, highlighted above as involved in actin cytoskeleton regulation, is an essential adaptor protein that promotes phagocytosis via activation of the small GTPase Rac and recruitment of the guanine nucleotide exchange factor *DOCK180 [85]. NCF1* codes for neutrophil cytosolic factor 1, a subunit protein of NADPH oxidase; NADPH oxidase produces reactive oxygen species that destroy the pathogen within the phagocytes [100].

It is also worth noting that intracellular bacteria enter the cytosol of their target cells by escaping from phagosomes during phagocytosis [87, 88, 98], or from endosomes during endocytosis [101]. Clathrin-mediated endocytosis is the main mechanism through which *P. salmonis* gains entry into the cytosol of Atlantic salmon macrophage cells [40], which are its main target cells [22]. In the current study, we detected differential methylation in 31 candidate genes that are involved in endocytosis such as *CLTA, EEA1, RAB5A* and *CBL-B. CLTA* encodes for the light chain of clathrin A. Clathrin is a structural membrane protein that has been reported to greatly aid the internalization of *P. salmonis* bacteria into Atlantic salmon macrophages [40]. Although we did not observe significant correlation between clathrin mRNA expression and methylation, a previous transcriptomic study of individuals from the same population reported differential expression of clathrin heavy chain and clathrin associated proteins in the head kidney [31]. *RAB5A* codes for the small GTPase Ras-related protein Rab-5A, which together with its effector early endosome antigen 1 (*EEA1*) promotes fusion of endosomes at early and late stages of endocytosis [102]. *CBL* codes for E3 ubiquitin-protein ligase CBL, involved in the ubiquitination of target proteins [103]. Interestingly, Moraleda et al reported a negative correlation between the expression of the E3 ubiquitin-protein ligase CBL gene and resistance to *P. salmonis* in Atlantic salmon [31]. Additionally, E3 ubiquitin-protein ligase *TRIP12* also showed differential methylation in the current study, further revealing potential involvement of protein ubiquitination and deubiquitination in P. *salmonis* infection. Indeed, several intracellular bacteria such as *Listeria ssp, Salmonella spp* and *Yersinia spp* have been reported to induce physiological changes in their target host cells by interfering with ubiquitination and deubiquitination processes to aid their survival and multiplication [103].

We also observed differential methylation of immune related genes between *P. salmonis* resistant and susceptible fish in the liver and head kidney. These genes and pathways are worthy of further investigation to elucidate the functional basis of genetic resistance to SRS in Atlantic salmon, and the potential role of methylation in this process. Nonetheless, while we observed important differences in methylation during the response to *P. salmonis* infection, only a few of these sites had significant associations with the mRNA expression of their annotated genes. Similarly, there was low overlap between differentially methylated sites and differential gene expression, a phenomenon also observed in previous similar study in Pacific salmon [41]. These observations may be attributed to the bulk nature of DNA methylation and transcriptome profiling, where cell diversity is not considered. The diverse cell populations in these tissues probably undergo unique DNA methylation and transcription alterations upon *P. salmonis* infection, lost via bulk tissue sequencing – this diversity can only be investigated via single cell transcriptome and DNA methylation sequencing. Additionally, there are multiple transcriptome regulatory mechanisms including histone modification, long and small non-coding RNAs, etc. that could play critical roles in modulating the Atlantic salmon response to *P.salmonis* infection, and it would be interesting to assess their involvement and their cross-talk with DNA methylation.

## 5. Conclusions

In the current study we investigated general DNA methylation patterns in the liver and head kidney of Atlantic salmon, as well as the methylation changes triggered by *P. salmonis* infection. Head kidney and liver present organ-specific methylation patterns, however the distribution of methylated sites across gene features and the methylation – gene expression trends are similar for both tissues. Although methylation was mostly negatively correlated with gene expression, there were a moderate number of positive correlations. Nonetheless, overall methylation towards the start and the end of the gene was associated with reduced expression. *P. salmonis* infection induced significant changes in the methylome of the head kidney, while the liver remained almost unaltered. These methylation changes regulated genes involved in crucial immune response pathways such as actin cytoskeleton regulation, pathogen recognition and phagocytosis. These results contribute to the growing knowledge on the immune response of Atlantic salmon to *P. salmonis* infection and may provide future avenues for the development of therapeutic strategies.

## Acknowledgements

The authors would like to thank the contribution of Benchmark Genetics Chile and Salmones Chaicas for providing the biological material and phenotypic records of the experimental challenges.

## Ethics declarations

The challenge experiment was carried out following the local and national regulatory systems and was approved by the Animal Bioethics Committee (ABC) of the Faculty of Veterinary and Animal Sciences of the University of Chile (Santiago, Chile), Certificate No. 01–2016, that based its approval decision on the Council for International Organizations of Medical Sciences (CIOMS)standards, in accordance with the Chilean standard NCh-324-2011.

## Consent to participate

Not applicable

## Consent to publish

Not applicable

## Data availability

All sequence data generated for this study have been submitted to the NCBI’s BioProject database under accession number PRJNA522369.

## Authors’ contributions

Conceptualization: DR, RDH, JMY; Formal analysis: RM, CP, AG, DR; Project administration: RDH, JMY; Writing - original draft: RM, DR, RDH; Writing - review & editing: All authors.

