## Supplementary File 1 for "The impact of *Piscirickettsia Salmonis* infection on genome-wide DNA methylation profile in Atlantic Salmon"

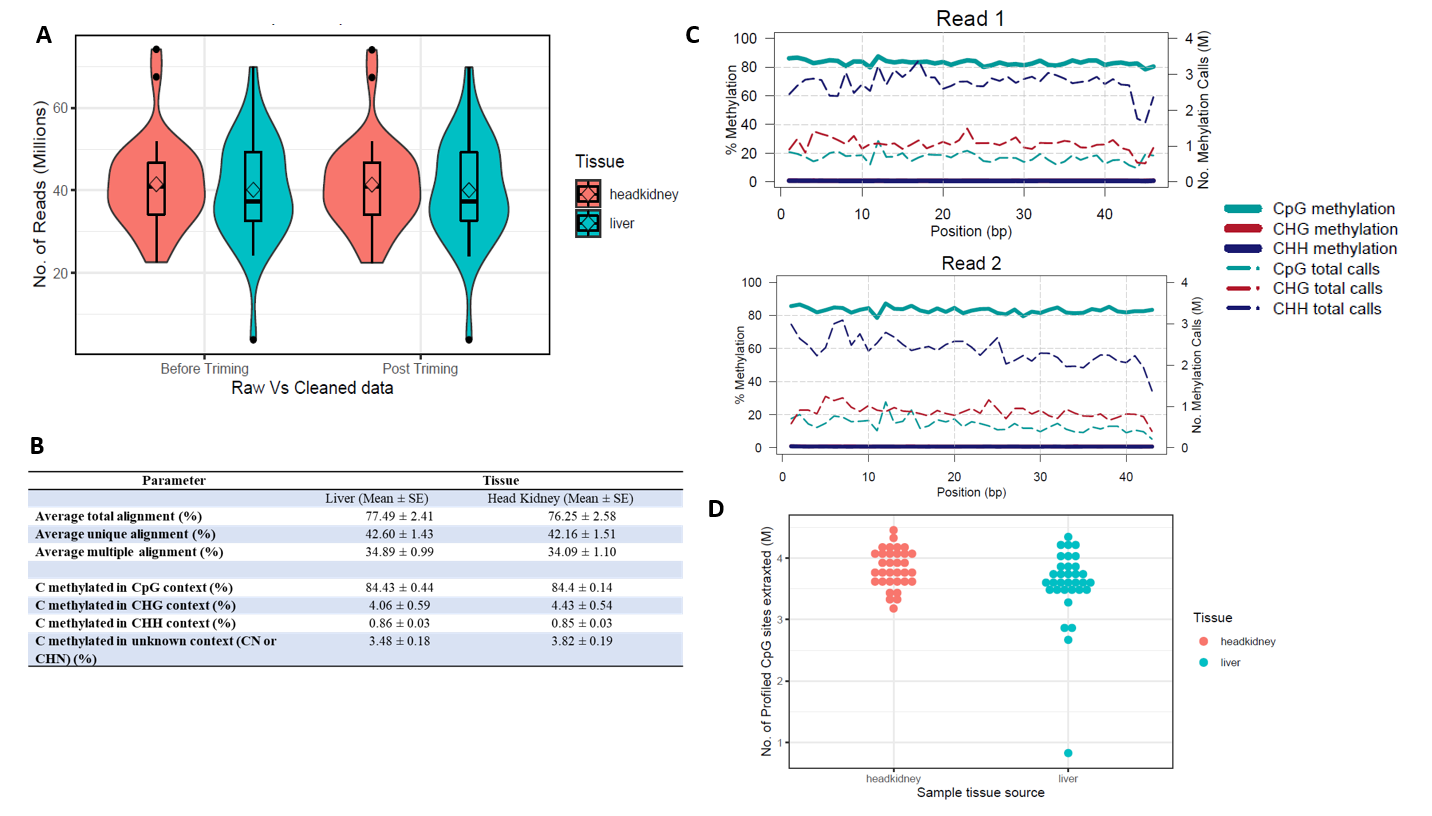

**Figure S1:** Sequencing and alignment quality; **A)** Violin plots showing sample variability in the number sequence reads from the RRBS library of each sample, **B)** Summary of alignment and methylation call quality, **C)** line plot showing different cytosine context calls and methylation along a sequence read pair, **D)** Violin plot showing the variation in the number of cytosine sites in the CpG context per sample.

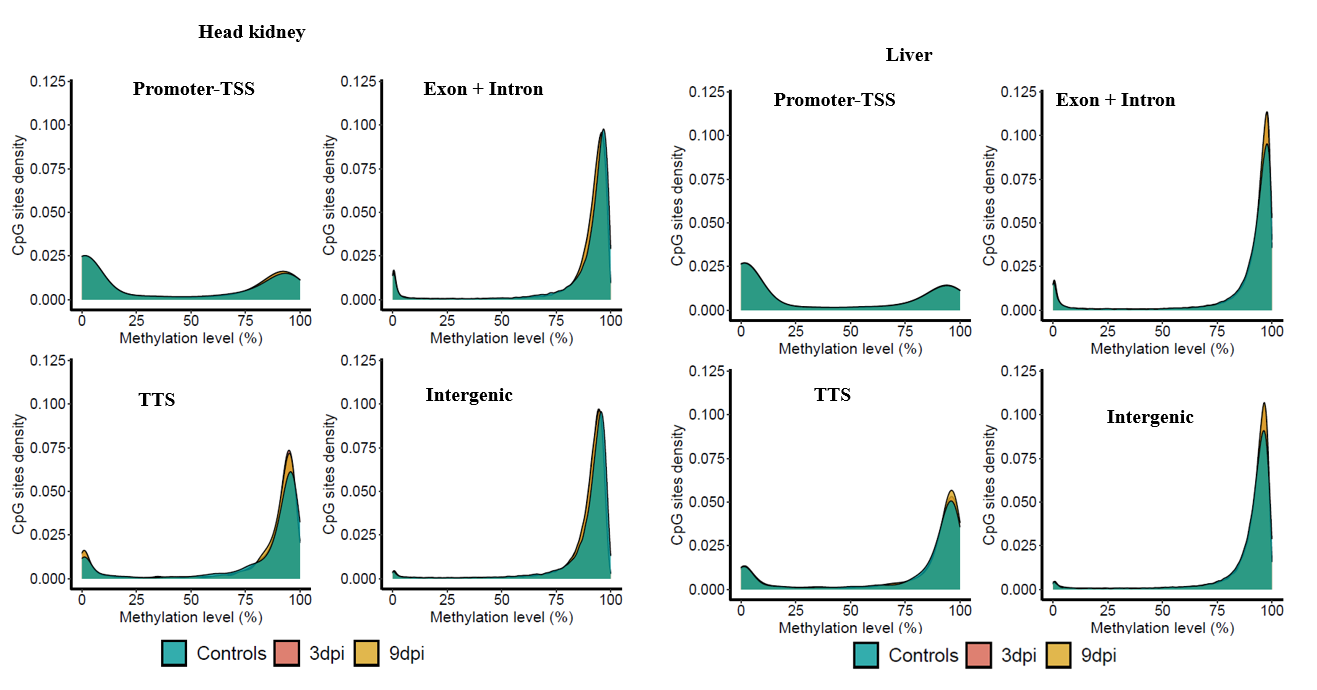

**Figure S2:** Density plots showing temporal methylation variability and distribution in the different genomic features for before and after infecting the fish with *P. salmonis* for head kidney and liver tissue.
